# Fast *in vivo* deep-tissue 3D imaging with selective-illumination NIR-II light-field microscopy and aberration-corrected implicit neural representation

**DOI:** 10.1101/2025.03.16.643569

**Authors:** Fenghe Zhong, Xue Li, Mian He, Yihang Huang, Chengqiang Yi, Shiqi Mao, Xin Huang, Kui Ren, Miaomiao Kang, Zhijun Zhang, Dong Wang, Dongyu Li, Peng Fei

## Abstract

Near-infrared II (NIR-II) microscopy, which enables *in vivo* deep-tissue visualization of vasculature and cell activities, has been a promising tool for understanding physiological mechanisms. However, the volumetric image speed of the NIR-II microscopy is hindered by scanning strategy, causing limitations for observing instantaneous biological dynamics in 3D space. Here, we developed a NIR-II light-field microscopy (LFM) based on selective-illumination and self-supervised implicit neural representation (INR)-reconstruction, which allows ultra-fast 3D *in vivo* imaging (100 volumes/s). Through integrating INR with view-wise aberration correction, we could conquer the artifacts induced by the angular subsampling and refractive index variation, achieving single-cell resolution at a volume of 550 μm diameter and 200 μm thickness. The volumetric selective-illumination overcomes the influence of out-of-focus background, together with the low scattering advantage of NIR-II wavelength, extending the imaging depth to 600 μm. The developed aberration-corrected implicit neural representation reconstruction (AIR) NIR-II LFM showcases its capability by monitoring hemodynamics of mouse brain under norepinephrine and flow redistribution of ischemic stroke in 3D vasoganglion, as well as noninvasively tracking immune cell activities inside subcutaneous solid tumor through intact skin. This approach represents a significant advancement in 3D *in vivo* imaging, holding great potential in biomedical research and preclinical studies.

## Introduction

*In vivo* optical microscopy has revolutionized biomedical research by enabling real-time visualization of cellular and even subcellular events in living animals, providing valuable insights on physiological mechanisms, disease pathogenesis, and therapeutic interventions. The continuous development of cutting-edge biomedical research demands that *in vivo* imaging technologies possess the capabilities of observing clearly, quickly, and deeply in 3D, posing a significant challenge due to the complex environment in the living body. A series of imaging modalities that could conquer the server tissue scattering have been intensively developed for understanding *in vivo* activities. Multiphoton microscopy achieved high-resolution at large depth using near-infrared (NIR) excitation light with less scattering and the corresponding point scan mechanism[1–3]. Photoacoustic microscopy, taking advantage of the much longer wavelength of ultrasound waves and its low transmission speed, achieves deep tissue volumetric mapping of vasculature in live animals[4–6]. As an emerging technique, NIR-II (900-1880 nm) fluorescence imaging explosives in biomedical research due to its long scattering length and remarkable penetration depth[7–10]. At the microscopic level, researchers realized optical sectioning from the cortical surface to 800 μm with long fluorescent wavelength at the NIR-II region and the confocal mechanism[11–13]. However, the above point-scanning-based acquisition strategies significantly degrade the temporal resolution, limiting their applications in highly dynamic life activities such as hemodynamics, especially for 3D monitoring.

Multiple approaches have been devised to reduce the scanning times to enhance the volumetric acquisition speed. Multi-focus two-photon microscopy splits a single excitation pulse into 30 spatially and temporally isolated pulses, enhancing the scanning speed by 30 times and achieving a volumetric imaging speed of 2 Hz[14]. By employing a spatial light modulator, temporal focusing technique excites multiple regions of interest simultaneously rather than sequentially, and captures images of different areas using one high-speed sCMOS camera, markedly boosting the image speed in 3D space[15]. Nevertheless, the severe scattering of visible light significantly compromises imaging depth. NIR-II light-sheet microscopy[16,17], which allows for selective-plane illumination and acquisition, achieves a high frame rate with large imaging depth owning to the long wavelength. However, its volumetric imaging speed remains constrained by the inherent scanning-based strategy.

Light-field microscopy (LFM), as a rising star in biological research[18–21], offers unique advantages for monitoring subcellular interactions and recording neuronal activities with high temporal resolution. By capturing projection images from multiple angles in a single shot, LFM facilitates volumetric reconstruction, achieving ultra-high temporal resolution 3D imaging[18,22]. Although some *in vivo* experiments have been demonstrated with LFM in visible wavelength, the very limited penetrating depth and the high requirements on sparse labeling impedes the practical use in biomedical research. Thus, the integration of NIR-II microscopy with LFM holds great potential for large-volume high-speed imaging of hemodynamics, tumor immunization, and neuronal activities. Firstly, the crosstalk fluorescence from out-of-focus areas poses a significant issue[23,24], particularly in thick organs such as the brain and tumors, where the fluorescence collected from beyond the focal zone can compromise accuracy and introduce artifacts. Additionally, besides the angular subsampling problem in LFM, which causes artefacts in 3D reconstruction[25,26], the limited angle range recorded with low NA objectives, often required for *in vivo* observations to achieve a sufficient field of view, exacerbates the missing cone problem[27–29] and further impacts reconstruction accuracy. Furthermore, non-uniform distributions of refractive index in deep-tissue generate spatially varying dynamic aberrations that impair imaging performance[29,30]. Aberration-induced degradation is particularly detrimental to LFM, as the cumulative effects of aberrations from multiple views aggregate. Although hardware-based adaptive optics have demonstrated efficacy in correcting aberrations in microscopic imaging[31,32], they are constrained by computational time and device refresh rates, resulting in correction times of several dozen seconds prior to each snapshot. This limits their applicability in high-speed LFM. Recently, a high-speed digital adaptive optics (DAO) algorithm tailored for LFM has been introduced[33]. This algorithm compensates for multi-angle aberrations through post-processing, thereby enhancing the reconstructed spatial resolution on *in vitro* cells and diverse tissue and organ surfaces. However, DAO simplifies the aberration model as linear translations in the light-field phase domain, with a solution space restricted to just two parameters. This approximation hinders its capability to address more intricate aberrations, thereby limiting its utility in deeper and more complex *in vivo* imaging contexts.

Here, we propose a NIR-II LFM for the first time, based on selective illumination and aberration corrected implicit neural representation (INR)-reconstruction, capable of deep-tissue volumetric imaging in living mice at video rate and cellular resolution. Complementary to the NIR-II LFM development, a bright NIR-II organic fluorescent nanoprobes with excellent photostability, designated as TTT8,4-B NPs, was also meticulously constructed. TTT8,4-B NPs help to enable volumetric imaging at an unprecedented speed of 100 Hz in living mice. To eliminate out-of-focus fluorescence and enhance the 3D reconstruction quality, we employ selective volumetric illumination achieved through rapid scanning of a light-sheet within the duration of a single snapshot. Instead of traditional deconvolution-based reconstruction methods, we developed an aberration-corrected implicit neural representation reconstruction (AIR) network for deep tissue imaging. Taking advantage of the continuity of the INR function and geometry consistency of synthesized views, our approach significantly reduces the impact of the angular subsampling problem and missing cone problem, and realizes higher axial resolution and fewer artifacts. Further integrated with a self-supervised view-wise aberration correction module in the network, our approach achieves minimized artifacts after reconstruction of a large volume (550 μm diameter and 200 μm thickness) with only 9 views and demonstrates significant performance advantage over deconvolution and cutting-edge DAO methods. We distinguished deep brain vasculature beyond the capability of a commercial two-photon microscopy with orders of reduced acquisition time. Our AIR NIR-II LFM facilitated the observation of 3D hemodynamics in the mouse brain due to its exceptional volumetric imaging speed and promising imaging quality, enabling us to analyze the response of the vasoganglion to clinical pressor norepinephrine and ischemic stroke. Furthermore, we functionalized TTT8,4-B NPs with specific antibodies to target immune cells *in vivo*, and utilized our imaging approach to achieve, for the first time, *in vivo* 3D tracking of immune cells in the tumor.

## Results

### Molecular design and photophysical properties

Motivated by the enhanced luminescence efficiency of nanoaggregates observed in fluorophores with aggregation-induced emission features, we focused on the design of bright NIR-II nanoprobes based on the aggregation-induced emission molecular scaffold. This boosted emission primarily arises from their characteristically distorted conformations, which increase intermolecular packing distances and mitigate non-radiative π–π stacking interactions, thereby promoting the retention of excited-state energy for radiative decay[34–38]. We, therefore, initiated our molecular engineering with TT6,2-B, a previously reported superior NIR-II aggregation-induced emission luminogen featuring a donor–π-bridge–acceptor–π-bridge–donor (D–π–A–π–D) architecture[39]. To further improve luminous brightness, we extended the π-bridge alkyl chains of TT6,2-B and incorporated additional triphenylethylene (TPE) rotors, successively obtaining TT8,4-B and TTT8,4-B (Fig. 1a and Fig. S1). These compounds were synthesized in high yields via classic reaction routes, with their structures and purity thoroughly confirmed (Fig. S2-S10). Photophysical characterization in tetrahydrofuran (THF) revealed their absorption peaks around 700 nm. Notably, the molar absorption coefficient (*ε*) increased steadily with both alkyl chain extension and the addition of TPE rotors, indicating enhanced NIR light-harvesting capability (Fig. S11). Owing to the inherently twisted structure, all compounds exhibited typical aggregation-induced emission peculiarity and broad NIR-II emission peaking at ∼1000 nm (Fig. S12).

**Fig. 1.**
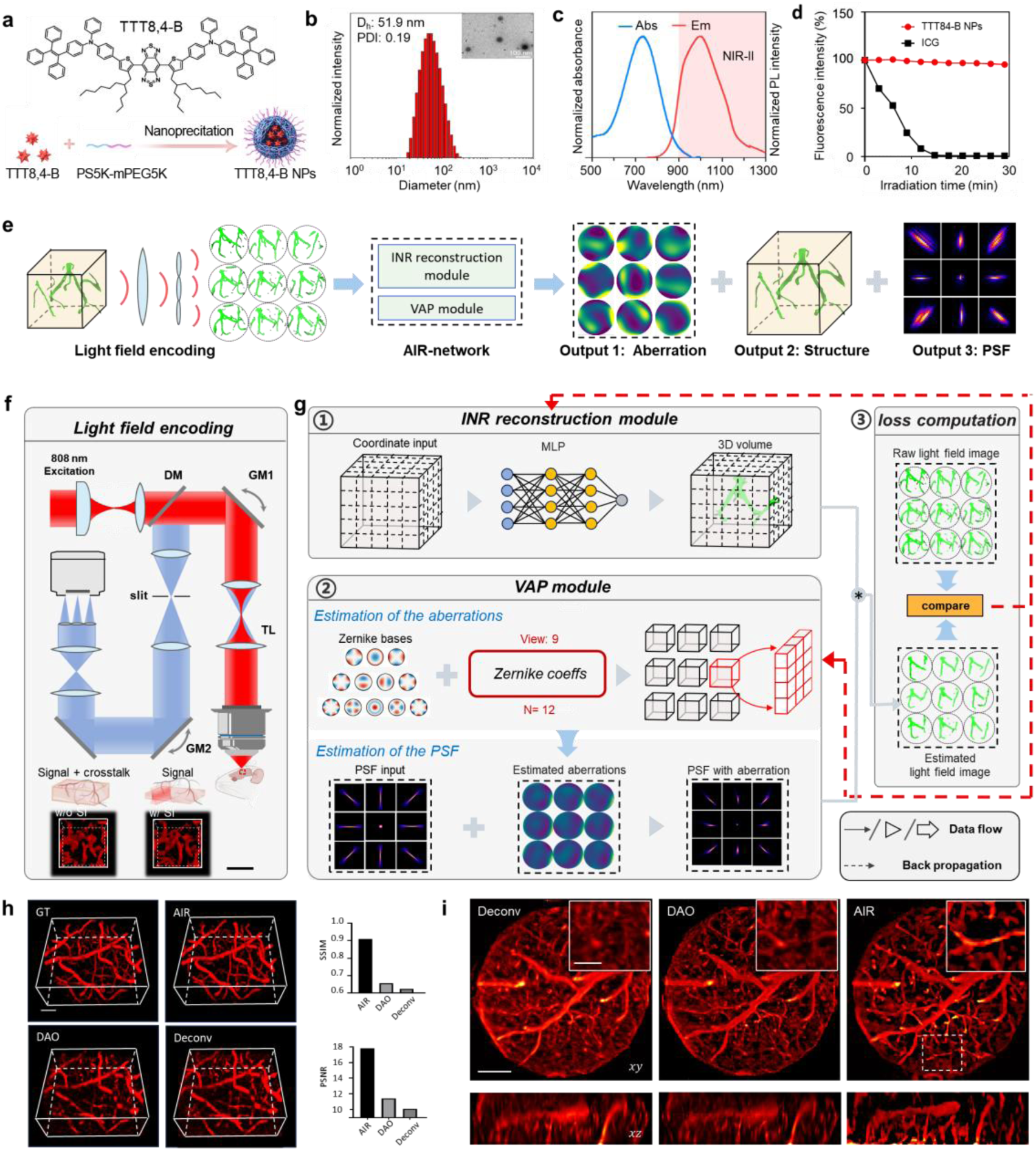
The NIR-II LFM based on selective illumination and Aberration-corrected INR Reconstruction (AIR). **a.** Chemical structure of TTT8,4-B and the fabrication of TTT8,4-B NPs. **b.** Hydrodynamic diameter (D_h_) and TEM image of TTT8,4-B NPs. **c.** Absorption and emission spectra of TTT8,4-B NPs. **d.** The photostability of TTT8,4-B NPs and ICG. **e.** The schematic of our AIR NIR-II LFM. **f.** Schematic of the light field encoding of selective illumination NIR-II LFM and the corresponding reconstruction. A cylindrical lens is employed to generate a light sheet, which is subsequently scanned by a galvanometer to enable selective-volume illumination. The excited fluorescence signal with limited crosstalk is modulated by microlens array and subsequently captured by a high-sensitivity camera. Scale bar: 100 μm. **g.** The algorithm decoding process of AIR which contains INR-reconstruction, VAP module and loss computation. **h.** The performance comparison of AIR, DAO and Deconvolution of a mouse brain vasculature simulated data, captured by two-photon microscopy. Scale bar: 100 μm. The SSIM of AIR, DAO and deconvolution reconstructed results are 0.91, 0.65 and 0.62. The PSNR of AIR, DAO and deconvolution reconstructed vasculature are 17.8, 11.4 and 10.1. Scale bar: 100 μm. **i.** The comparison of reconstructions by deconvolution, DAO and AIR. The white boxed zoom in (right) indicates the noticeable errors in deconvolution and DAO based reconstruction result which is accurately resolved by the AIR method. Scale bar: 100 μm (left, right).

In order to facilitate the *in vivo* application, these hydrophobic compounds were formulated into water-dispersible nanoparticles (NPs) *via* the nanoprecipitation method[40] using amphiphilic copolymers (PS5000-mPEG5000). Dynamic light scattering test revealed that the hydrodynamic diameters (D_h_) of TT6,2-B, TT8,4-B, and TTT8,4-B NPs were approximately 92, 56, and 52 nm, with transmission electron microscopy (TEM) confirming their spherical morphology (Fig. 1b and Fig. S13). The zeta potentials of these NPs were determined to be -20.9, -16.7, and -19.7 mV, respectively (Fig. S14). These NPs showed a significant red-shift in absorption maximum to ∼ 730 nm compared to that in their THF solution, benefiting long-wavelength excitation and improved tissue penetration depth, and exhibited strong fluorescence in the 800–1300 nm range, peaking at 1000 nm, indicating their potential as excellent NIR-II imaging probes (Fig. 1c and Fig. S15). Thus, using a robust NIR-II reference dye with a known absolute quantum yield (QY) from our previous work[41], we determined the relative QYs of TT6,2-B, TT8,4-B, and TTT8,4-B NPs in aqueous solution (900–1400 nm) to be 0.57%, 0.69%, and 0.80%, respectively (Fig. S16). In contrast, the relative QYs of two commercial NIR-II dyes including indocyanine green (ICG) in water and IR-26 in 1,2-dichloroethane were measured to be only 0.19% and 0.04%, respectively (Fig. S17), which are approximately 4.2- and 20-fold lower than that of our best-performing probe (TTT8,4-B NPs). These results demonstrated that the introduction of longer alkyl chains and more rotors enhanced the fluorescence brightness of molecular aggregates. Furthermore, TTT8,4-B NPs exhibited excellent photostability (Fig. 1d and Fig. S18) and remarkable storage stability (Fig. S19), making them a superior candidate for NIR-II fluorescent imaging.

Systematic theoretical simulations and computational analyses were conducted to understand molecular design principles. Initially, the ground-state (S_0_) geometries of these three compounds confirmed their extremely twisted configurations (Fig. S20). Moreover, insignificant variation in the energy gaps between the highest occupied molecular orbital (HOMO) and the lowest unoccupied molecular orbital (LUMO) for these compounds was recorded, spanning from 1.349 to 1.426 eV (Fig. S21), supporting their almost identical absorption and emission wavelengths. Subsequently, TT6,2-B and TTT8,4-B were selected to investigate the influence of alkyl chain and additional rotor on molecular packing in aggregates by molecular dynamics simulations (Fig. S22-S23). As depicted in Fig. S24-S26, due to the incorporation of TPE units and the elongation of alkyl chains, TTT8,4-B exhibited higher values of solvent-accessible surface area and radius of gyration (Rg) than that of TT6,2-B, demonstrating its looser packing structure. Additionally, TTT8,4-B aggregates form fewer hydrogen bonds with water, highlighting its stronger hydrophobic character, thus reducing the twisted intramolecular charge transfer (TICT)-caused fluorescence quenching in high-polarity, aqueous physiological environments[42]. Together, the decreased intermolecular π-π stacking interaction and TICT effect promote the radiative transition and elevated QY of TTT8,4-B in nanoaggregates.

The biosecurity of TTT8,4-B NPs was thoroughly evaluated, showing excellent biocompatibility *in vitro* through cytotoxicity assessments and hemolysis assays (Fig. S27). *In vivo* studies *via* tail vein injection of TTT8,4-B NPs revealed obvious signal accumulation firstly in the liver at 24 h post-injection, decreasing significantly over time (Fig. S28). Assessments of physiological parameters, including body weight, liver and renal function markers, and blood routine parameters, alongside H&E staining of major organs, indicated no significant differences between TTT8,4-B NPs and PBS-treated mice (Fig. S29-S31 and Table S1). Overall, these results demonstrate that TTT8,4-B NPs are a promising candidate for high-performance NIR-II fluorescence imaging *in vivo*.

### The principle of AIR selective-illumination NIR-II LFM

Interrogating the fast 3D dynamics in live animals is crucial for revealing biological mechanisms[43–45]. However, such complex imaging condition brought by light scattering and attenuation rises the challenges on the volumetric imaging speed, imaging depth and anti-aberration ability of microscopes. These metrics entangled together manifested as the tradeoff of imaging quality and imaging speed in scanning-based microscope[46,47].

Here, to capture the fast 3D time-varied signals, we proposed a NIR-II light field microscopy with selective volume illumination and aberration-corrected INR reconstruction algorithm to divide-and-conquered this problem by optimizing the signal generation, optical encoding, and algorithm’s decoding.

To efficiently collect the 3D information in living animals in a snapshot, one plausible approach lies in the development of light-field microscopy[48]. However, this method alone suffered from the cross-talk due to the out-focus signals in thick tissue. Sequential scanning-based approach[49] could solve this problem but sacrificed the volumetric imaging speed severely. Taking both into consideration, we have developed the selective illumination-armed Fourier light field microscopy (FLFM, Light field encoding in Fig. 1e, Fig. 1f and Fig. S32). This innovative technique rapidly scans the imaging volume with light-sheet excitation in one light-field imaging duration, thereby achieving both the high-efficiency volumetric encoding and out-of-focus crosstalk minimization. This improved excitation strategy benefited the signal to background ratio (SBR) and weak signal detection (Fig. S33). To generate the selective volume illumination, a cylindrical lens is employed to generate a light sheet, which is subsequently scanned by a galvanometer. The fluorescence light is collected by the objective lens and descanned by the same galvanometer as the illumination beam, purified the in-depth signal, scanned and split by a microlens array. With the aim of observing living organisms at single-cell resolution, we customized a microlens array with square arrangement to fully split the optical aperture into 9 FLFM views, which has an effective NA of 0.14 for each microlens (Methods), producing 4.27 μm × 9.5 μm spatial resolution and ∼200 μm depth of field (DOF), and enables capturing the vasculature in mouse ear, iris, brain, subcutaneous tumor (Fig. S34) and neuronal dynamics after sciatic nerve injury (Fig. S35). The Rayleigh range of the light sheet was designed as 69 μm to maximal the remove of the crosstalk out of the 200 μm DOF while maintaining an acceptable signal amplitude in the region of interest. We characterized NIR-II LFM by imaging fluorescent beads embedded in agarose gel. The system can achieve a spatial resolution of 4.7 μm × 11 μm (Fig. S36).

After acquiring the enhanced light-field captures, accurately decoding the 3D signals from the encoded 2D multi-view projections remains a significant challenge for existing light-field reconstruction algorithms[18]. Besides of the information compression in light-field encoding, which cause a severer resolution degradation, the presence of optical aberrations, derived from the dynamic refractive index inhomogeneities in *in vivo* deep-tissue imaging, further complicates the 3D reconstruction inverse problem, manifested as the view-wise wavefront distortion[33].

While supervised learning approaches[18] would achieve resolution improvement, they suffered the generalization errors and struggle to build an accurate degradation model especially under this unknown complex aberration (varied with time and view-angle). In contrast, classic-model based approach[33] mitigated the aberration without prior knowledge, but remained limited by their simplified aberration models.

Here, we employ an AIR network, which contains an INR reconstruction module and a view-wise Aberration prediction (VAP) module, to reduce the artifacts raised by sparse angular sampling rate and aberrations in *in vivo* imaging (AIR-network in Fig. 1e and Fig. 1g). First, a 3D volume was generated with coordinates input during Multi-Layer Perceptron (MLP) inference. The wave-optics model was then introduced to project the 3D volume onto a 2D estimated light-field map which act as a constrain. To obtain an accurate forward model, the VAP module incorporates learnable Zernike coefficients into light-field PSF to estimate the view-wise aberrations (Supplementary Text S1). The estimated phase of different views in Fig. 1g described the wavefront distortion from objective space (biological sample) to imaging system (FLFM system) in an end-to-end manner, achieving higher fidelity of PSF estimation (Fig. S37) and 3D reconstruction than conventional aberration-correction approach (Fig. S38). By calculating the loss between the LF projections with estimated aberration and the raw LF captures, the INR network could gradually remove the artifacts induced by aberrations in 3D reconstruction (Fig. S39). Through offline processing, our method could estimate the phase of each FLFM view and produce accurate reconstructions during *in vivo* FLFM imaging (Fig. 1g). Notably, our approach does not rely on hardware correction devices or additional prior information, offering a highly efficient and accessible solution for aberration correction and 3D reconstruction especially in acute biological dynamics.

Mouse brain vasculature, captured by two-photon microscopy, was first used to validate the accuracy of AIR reconstruction. We projected the volumetric vasculature in the nine views corresponding to our MLA setting. As shown in Fig. 1h, the reconstructions by AIR, DAO, and deconvolution were performed and compared with the ground truth. The SSIM of AIR, DAO, and deconvolution reconstructed results are 0.91, 0.65, and 0.62. The PSNR of AIR, DAO, and deconvolution reconstructed vasculature are 17.8, 11.4, and 10.1. These two factors both indicate AIR’s excellent performance and high accuracy of the mouse brain vasculature reconstruction with only 9 views.

We then demonstrated the performance advances of AIR through a comparison with DAO and conventional deconvolution-based reconstruction method on *in vivo* mouse brain vasculature. As shown in Fig. 1i and Movie S1, in deconvolution-based reconstruction image, the reconstructed vessels have severe artifacts, as well as elongations along the axis. When DAO is used in FLFM, it can alleviate the problem of artifacts to a certain extent. However, there are still vascular discontinuities due to signal loss. Also, the DAO does not solve the axial elongation caused by the missing cone problem. In sharp contrast, AIR reconstructs more details both horizontally and axially. Furthermore, due to the intrinsic advantage of continuity of the INR function and geometry consistency of synthesized views, our method reduces the axial elongation while reconstructing 3D image. The versatility of AIR has also been validated with a thick GFP-labeled brain slice which captured by a visible light-field microscopy system (Fig. S40) and a consistent result has been observed.

### NIR-II and visible LFM comparison in phantom

To showcase the capability of our AIR NIR-II LFM in obtaining high-resolution volumetric images in deep tissue, we prepared leaf phantom immersed in lipids, mimicking the tissue environment (Fig. 2a). The phantom was stained with a mixture of Rhodamine B and TTT8,4-B NPs solution. Imaging of the phantom was conducted using both visible LFM (532 nm) and NIR-II LFM, through lipid agarose gel layers with varying thicknesses. The results reveal that NIR-II LFM can clearly resolve the intricate leaf vasculature from the surface down to a depth of 500 μm under lipids (Fig. 2b). In contrast, the quality of vasculature reconstruction in visible LFM significantly deteriorates with the addition of just a 100 μm lipid layer, and the vessels became indistinguishable with artifacts when the lipid layer reaches 200 μm (Fig. 2c). We further quantified the structure similarity index measurement (SSIM) between the surface MIP images and the deeper layer images (Fig. 2d). Compared to visible group, NIR-II group exhibits consistent SSIM values, highlighting the deep penetration and reconstruction capabilities of NIR-II LFM in scattering tissue. Conversely, the SSIM of images captured by visible LFM decreases markedly.

**Fig. 2.**
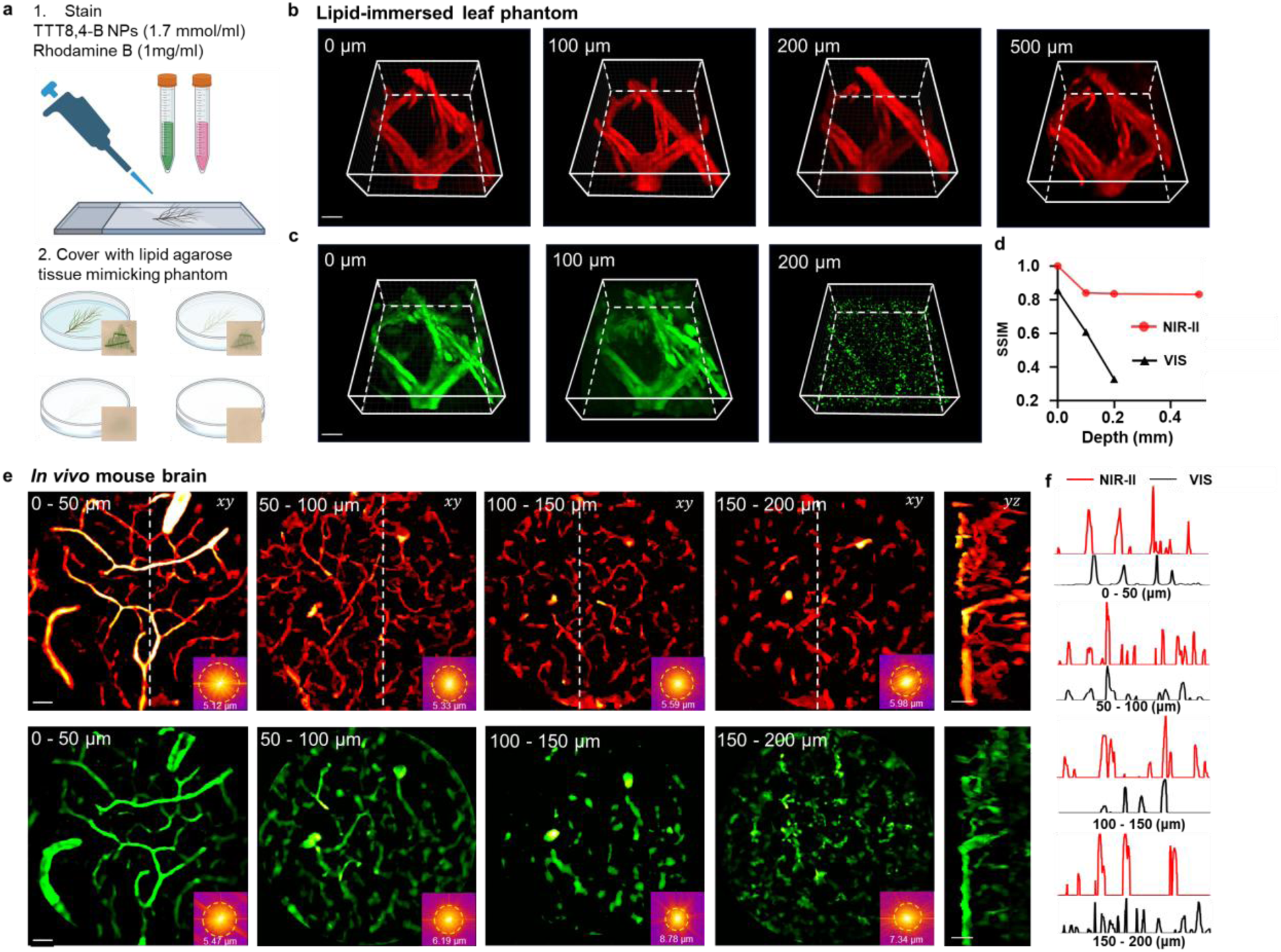
Visible and NIR-II LFM comparison on leaf phantom and *in vivo* mouse brain. **a.** The procedure of leaf phantom preparation. **b.** TTT8,4-B NPs and Rhodamine B co-dyed leaf vasculature imaging with NIR-II LFM. Scale bar: 50 μm. **c.** The leaf vasculature imaging with the visible LFM. Scale bar: 50 μm. **d.** The degradation of SSIM while imaging with different depths of lipid phantom covered. **e.** Visible and NIR-II co-registered *in vivo* mouse brain imaging. Scale bar: 50 μm. **f.** The cross-sectional profiles of the dashed white lines in **e**.

### NIR-II and visible LFM *in vivo* mouse brain comparison

The unique advantage of our NIR-II LFM over traditional visible LFM was further validated with *in vivo* mouse brain imaging. We performed a side-by-side comparison between the two imaging modalities in the same region. As shown in Fig. 2e, the NIR-II LFM achieved high imaging quality with little resolution loss at different depths (spectrograms in the right bottom), while the resolution of visible microscopy degraded significantly from 5.5 μm to 8.8 μm. Cross-sectional profiles in Fig. 2f show that there are microvessel missing using the visible LFM due to the short scattering length of visible light and the low signal-to-background ratio (SBR) of the acquired images.

### Deep tissue hemodynamic imaging

Fig. 3a presents a mosaic of partially overlapping reconstruction volumes, seamlessly integrating to form a continuous vasculature map that rivals the imaging depth capabilities of other scanning-based imaging techniques[50]. The vasculature is distinctly resolved, spanning from the cortical surface down to a depth of 600 μm. Fig. 3b illustrates the vessels at three distinct depths (surface, 300 μm, and 600 μm), with the depth sectioning further demonstrated in Movie S2. Dyes with longer emission spectrum (nanocrystals[51], 1500 nm), can further extend the penetration depth to 800 μm below the cortex surface. As shown in Fig. S41, our AIR-enabled NIR-II light field microscopy (LFM) achieves a volumetric acquisition speed nearly 100 times faster than conventional two-photon microscopy (2 Hz vs. 0.02 Hz for ∼50,000,000 μm³) at an imaging depth of 800 μm, and previously indistinguishable and blurred vasculature captured by two-photon fluorescence microscopy (Olympus FVMPE-RS) can be clearly resolved using our approach.

**Fig. 3.**
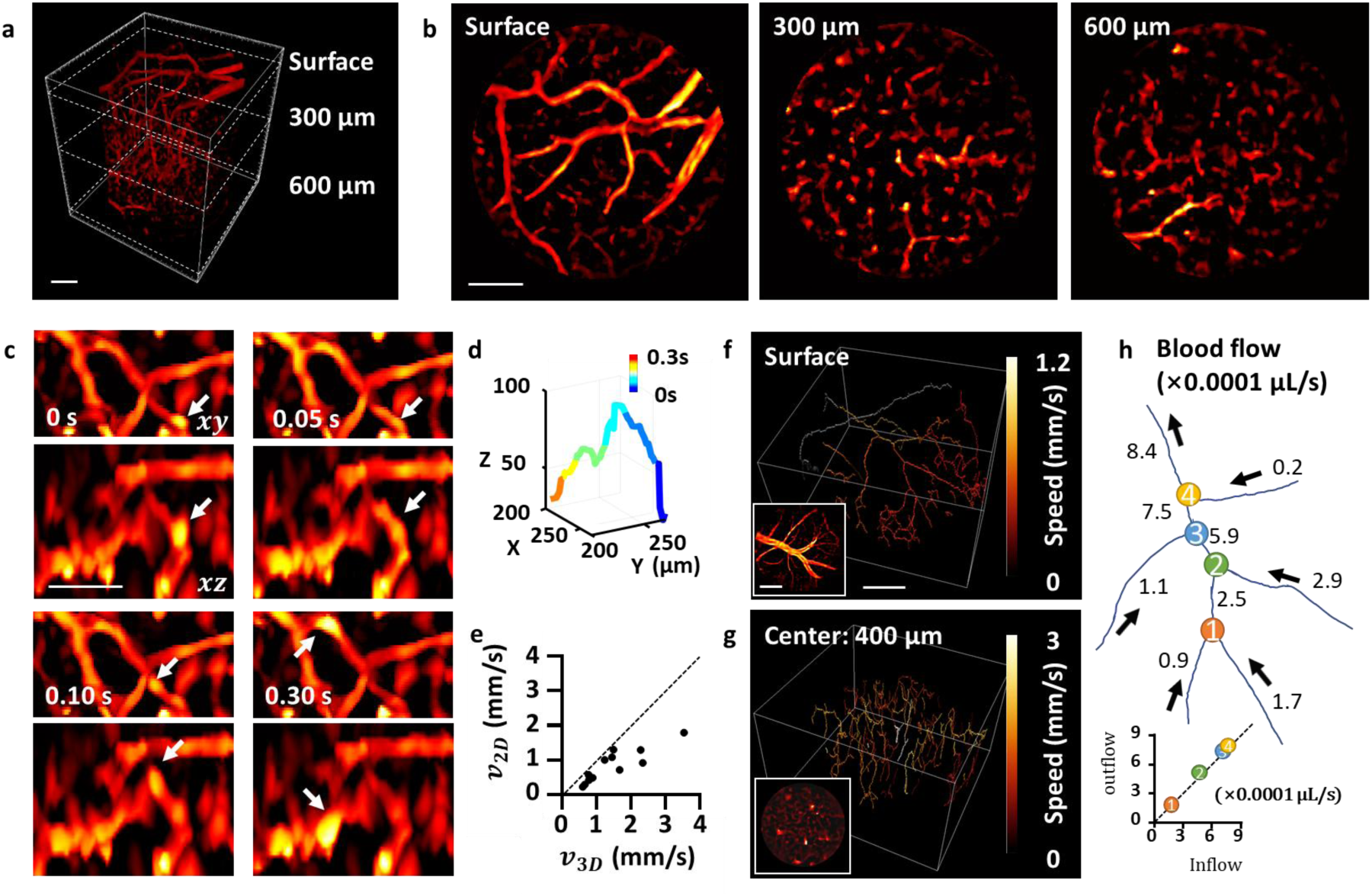
The performance of AIR NIR-II LFM. **a.** The volume rendering of the *in vivo* mouse brain vasculature from the cortical surface to 600 μm. Scale bar: 100 μm. **b.** The vasculature at the motor cortex surface, 300 μm and 600 μm depth. Scale bar: 100 μm. **c.** Time-lapse *xy* and *xz* MIP images of the vessel with the flowing particle cluster outlined by the white arrow. Scale bar: 50 μm. **d.** The time-lapse 3D path of the flowing particle cluster. **e.** Flow velocity comparison between 2D (*v*_2*D*_) and 3D (*v*_3*D*_) quantification methods. **f-g.** Color-encoded flow velocity in the mouse brain skeleton (surface and 400 μm in depth). Scale bar: 100 μm. **h.** Vessel segmentation reveals dynamic changes in **f** blood flow and the conservation of the volumetric inflow and outflow rates at bifurcation.

Non-invasive capture of hemodynamics and blood flow *in vivo* offers profound insights into neuroscience and cancer studies that highly relate to metabolism. Our AIR NIR-II LFM, as a high temporal-resolution volumetric imaging tool, presents an accurate method for blood flow quantification in 3D spaces. Fig. 3c showcases a region within the volume image, with the white arrow highlighting the flowing particle cluster of interest (probably TTT8,4-B NPs conjugated with hemoglobin or other biomacromolecules in blood). These MIP images clearly illustrate volumetric movement in both lateral and axial directions. Notably, the high reconstruction quality allows for the analysis of each vessel branch’s depth and penetrating direction. The time-lapse 3D path of the flowing target could be extracted and presented in Fig. 3d. The incorporation of depth information enhances the accuracy of flow velocity analysis compared to conventional 2D kymograph analysis methods[52]. The differences in the flow velocities quantified by these two methods are illustrated in Fig. 3e. Despite the depth, high imaging frame rate is also essential for accurate flow speed quantification. Operating such a high volumetric imaging rate at 100 Hz (550 μm diameter and 200 μm thickness), flowing particle clusters can be resolved by tracking successive volumes (Movie S3). The flow velocities of the selected vessels were quantified and shown in the color-encoded vessel skeleton images in Fig. 3f (Surface) and Fig. 3g (Center depth: 400 μm). Combining the high-resolution vasculature and blood flow speed of each vessel segment between the two adjacent bifurcations, we further computed the volumetric blood flow of each segment in absolute values (Fig. 3h). The inflow and outflow rates show a conserved blood flow volume, which further verified our AIR NIR-II LFM’s quantitative anatomical and flow measurement ability.

### Mouse brain *in vivo* imaging under norepinephrine

To demonstrate the capability in capturing acute brain activities, norepinephrine, which is used to increase the artery pressure in the clinic, was injected while imaging (Fig. 4a). The contraction in the artery was clearly observed (Fig. 4b). Time-lapsed cross-sectional profiles of the selected artery (yellow arrow heads in Fig. 4b) and vein (blue arrow heads in Fig. 4b) are shown in Fig. 4c. After norepinephrine injection, the artery underwent contraction, reducing its volume to 60% of its resting state. The vein exhibited a more limited reduction in size (∼10%). Additionally, vessel contraction was evident along the axial direction. The dynamic response of the vessel, captured with high temporal resolution, is presented in Movie S4. We further quantified the corresponding flow velocity, as depicted in Fig. 4d, revealing a significant increase. The concomitant occurrence of vessel contraction and elevated flow velocity suggests an increase in blood pressure within the brain vessels, aligning with clinical prediction.

**Fig. 4.**
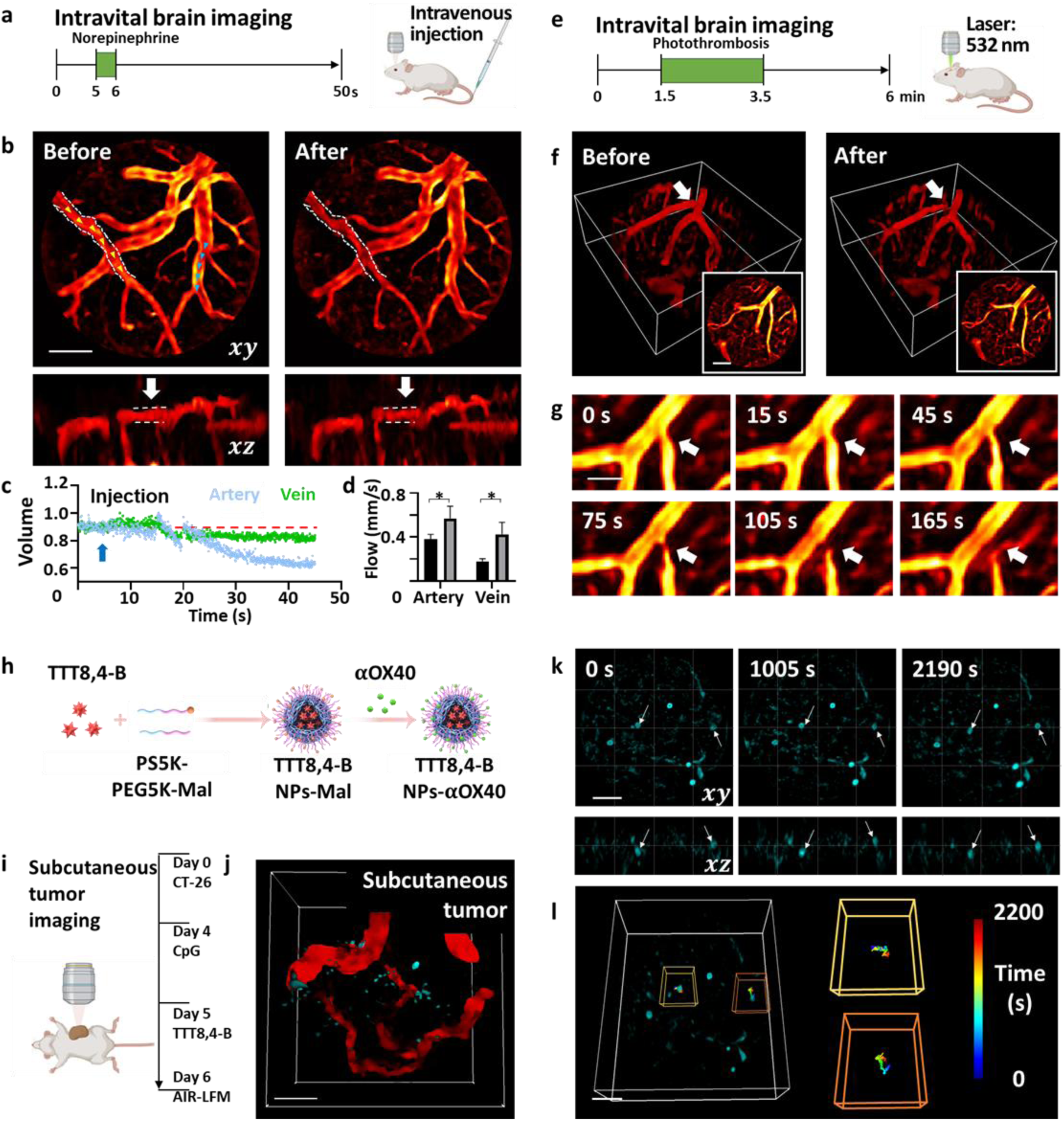
Mouse brain hemodynamic monitoring and T-cell tracking in a solid tumor. **a.** Schematic of norepinephrine experiment. **b.** MIP of the vasculature before and after norepinephrine injection. Left: before injection, right: after injection. Top: *xy* projection, bottom: *xz* projection of the selected region. The dashed lines indicate the vessel diameter changes before and after injection. Scale bar: 100 μm. **c.** The volumetric change of the target artery and vein during the experiment. The blue arrow indicates the time of injection. The red line indicates the normalized diameter before the injection. **d.** The flow velocity comparison before and after the norepinephrine injection. **e.** Schematic of ischemic stroke experiment. **f.** The volume renderings and the *xy* projected images before and after the ischemic stroke. Scale bar: 100 μm. **g.** Time-lapse monitoring of the generation of ischemic stroke and flow redistribution. Scale bar: 50 μm. **h.** A schematic illustration for preparation of the TTT8,4-B NPs-αOX40. **i.** Schematic of the tumor experiment. **j.** Dual-color imaging of vasculature and T cells with AIR NIR-II LFM in subcutaneous tumor. Scale bar: 100 μm. **k.** T cells monitoring in subcutaneous a tumor. The white arrows indicate the three moving cells. Scale bar: 100 μm. **l.** T cell volumetric tracking in a subcutaneous solid tumor. Scale bar: 100 μm.

### Monitoring the formation of ischemic stroke

Ischemic stroke is the most common type of stroke, which can cause permanent brain damage and be fatal. Monitoring the formation of clots with high spatial-temporal resolution presents a unique opportunity for the development of potential therapies for ischemic stroke.

As shown in Fig. 4e, artery-targeted photothrombosis was applied while monitoring with the AIR NIR-II LFM. Prior to the experiments, Rose Bengal and TTT8,4-B NPs were administered. Fig. 4f presents the vasculature before and after stroke induction. The targeted artery was occluded using a 532 nm laser for excitation. The subsequent clot formation, accompanying changes in vessel morphology, and 3D flow redistribution were observed and illustrated in Fig. 4g and Movie S5.

### Noninvasive *in vivo* imaging of T cells in subcutaneous solid tumor through intact skin

The high penetration depth and high temporal-spatial resolution of the AIR NIR-II LFM enable noninvasive 3D *in vivo* imaging and real-time tracking of T cells in the subcutaneous solid tumor through intact skin. For ease of observation, cytosine-phosphate-guanine (CpG) was intratumorally injected to activate T cells previously because it is the ligand of Toll-like receptor 9 (TLR9) and is capable of inducing systemic immune responses upon binding with TLR9[53,54]. To facilitate the targeted monitoring of T cells within the tumor microenvironment, TTT8,4-B NPs were functionalized with αOX40 to yield TTT8,4-B NPs-αOX40 (Fig. 4h). αOX40 is the monoclonal antibody of OX40, which is a co-stimulatory molecule expressed on activated effector T cells and regulatory T cells[55,56].

With the assistance of T cells-targeted TTT8,4-B NPs-αOX40, vessel - T cells dual-color imaging can be pictured in subcutaneous tumor environment (Fig. 4j). More importantly, the movements of the T cells were imaged by the NIR-II LFM with 500 ms integration time each 15 s (Movie S6) following the preparation in Fig. 4i. In this case, we observed the cell movements and mapped its trajectories. As shown in Fig. 4k, a location shift during 30 min can be observed volumetrically. Fig. 4l shows the time-lapsed trajectories of the three target T cells. Such results suggest our approach holds great potential in the investigation of immunotherapy of solid tumors.

## Discussion

Observation of biological dynamics *in vivo* in deep tissue is essential for investigating physiological mechanisms, disease pathogenesis, and therapeutic interventions. However, it is challenging due to the inherent trade-off among visualizing clearly, deeply, and quickly. Scanning-based microscopy offers high resolution and large penetration depth, but severely sacrifices the imaging speed, thereby limiting the capability of capturing fast biological dynamics in 3D spaces. Here, we introduce a NIR-II light-field microscopy technique capable of high-quality, deep-tissue volumetric imaging at an unprecedented temporal resolution. It is worth noting that such an efficient combination is based on a series of technical advances. First, to address the challenge of poor 3D reconstruction quality stemming from sparse and limited angular captures, particularly pronounced in large-scale *in vivo* observations due to the low NA of the objective, we introduce physics-based INR for 3D reconstruction. Leveraging the continuity of the INR function and geometry consistency of synthesized views, our approach significantly reduces artefacts and improve axial resolution. More importantly, our VAP module, the self-supervised digital AO, view-wisely corrects the abbreviations introduced by both the optical system and complex samples, without compromising capturing speed as hardware AO does, allowing LFM to fully exploit its advantage of high temporal resolution. The AIR reconstruction performed on visible and NIR-II LFM data both showcase the excellent performance over other digital based aberration correction method (DAO). Furthermore, it is crucial to prevent out-of-focus fluorescence for LFM reconstruction in thick tissues. Our scanning lightsheet-based volumetric selective illumination approach significantly enhances imaging quality and facilitates the differentiation of small structures that would otherwise be obscured without selective illumination.

While NIR-II LFM exhibits enhanced performance over existing *in vivo* optical microscopy techniques, particular in terms of 3D imaging speed, it still faces limitations. TTT8,4-B NPs emits fluorescence with a peak located at 1000 nm, although belongs to NIR-II region, it still suffers from relatively higher tissue scattering compared to longer wavelengths. The development of bright, biocompatible, multi-color, and targeted fluorophores with longer emission wavelengths would boost detection sensitivity at depth, enhance the capacity of this detection technology.

## Conclusions

Our newly developed NIR-II LFM strategy circumvents the compromise between speed, resolution, and depth in *in vivo* imaging based on a fully self-supervised view-wise aberration corrected INR reconstruction and volumetric selected illumination.

With the integration of ultra-fast LFM, customized AIE NPs, and deep NIR-II detection, our strategy provides a volumetric imaging speed of 100 Hz for a volume of 550 μm diameter and 200 μm thickness at cellular resolution and finally achieves a large imaging depth of 600 μm in the mouse brain, while visible LFM could not obtain distinguishable vessels over only 100 μm depth. Our approach also allowed to capture the artery contraction and vein dilation under the challenge of norepinephrine, as well as to monitor hemodynamic redistribution during the formation of ischemic stroke through photothrombosis. Moreover, the 3D movement of immune cells inside a subcutaneous tumor was clearly observed through intact skin and trajectory tracking could be easily performed. Therefore, AIR selective-illumination NIR-II LFM can be a powerful tool in brain science and tumor science. Potential investigations of neuronal activity and tumor environment would further elevate the impact of this technique.

## Methods

### NIR-II LFM

A schematic diagram of the experimental arrangement is shown in Fig. S32a. The laser beam from a NIR CW laser (808 nm, 200 mw, Changchun New Industries Optoelectronics Tech, Co., Ltd.) is collimated with an objective lens. A beam expander, which is constructed by two lenses (L1, f = 45 mm, OLD2435-T3; L2, f = 60 mm, OLD2438-T3 JCOPTIX), is used to reshape the beam diameter. A cylindrical lens (CL1, f = 200 mm, OLYC2601661-T3, JCOPTIX) and multiple doublets (L3, f = 30 mm, OLD2430-T3; L4, f = 60 mm, OLD2438-T3; L5, f = 100 mm, OLD2444-T3, JCOPTIX) are selected to adjust the beam waist in two directions. The NIR beam is steered by a high-speed galvanometer (6240H, Cambridge Technology, INC.) and passes through a dichroic mirror (DM1, OFD1LP-950, JCOPTIX). A doublet (L6, f = 250 mm) and a tube lens (TL, f = 200 mm, ACT508-200-C, Thorlabs) relay the galvanometer to the back aperture of the objective lens (25x, 1.0NA, XLPLN25XSVMP2, Olympus). The generated fluorescent light is collected by the objective and then reshaped by the TL and lens (L7, f = 200 mm, ACT508-200-C, Thorlabs). A micro lens array (pitch size 2.4 mm, f = 30 mm) is carefully adjusted to generate view images from 9 directions. In this study, we enhance the imaging depth-of-field without compromising the field-of-view by designing a mask for each microlens to reduce their light-gathering aperture (d = 1.7 mm). The images are collected by relay lens L8 (f = 75 mm, OLD3240-T3, JCOPTIX) and L9 (f = 75 mm, OLD3240-T3, JCOPTIX) on camera (pixel = 15 um, Ninox Ultra 640, Raptor Photonics). Another CW laser at 532 nm (MGL-U-532-20mW, Changchun New Industries Optoelectronics Tech, Co., Ltd.) is coupled into the system by the DM1 to generate a tightly focused spot on the mouse brain for photothrombosis. For deep tissue imaging, the system is slightly adjusted and aligned in Fig. S32b described in the selective illumination section.

### Selective illumination

Fluorescence microscopy suffers from the out-of-focus background signals due to the linearity of single photon absorption. This image quality degradation, especially on thick tissue, casts doubt on the accuracy of light field reconstruction. Two-photon scanning LFM[49] eliminates crosstalk by two-photon absorption, but it is time consuming and the ultrafast system is relatively expensive. Confocal alignment with a specially designed mask[44] shows excellent performance but also places high requirements on design, fabrication and alignment.

To improve the reconstruction quality, out-of-focus crosstalk was mitigated through carefully designed selective illumination. As shown in Fig. 1a and Fig. S32b, the thin illumination plane is generated by a cylindrically focused lens. By forming a focused light-sheet plane and scanning with a galvanometer, only the on-focus reconstruction region was illuminated with high fluence. The beam waist of the light sheet is designed to 3 μm which corresponds to a Rayleigh range of 69 μm. Then, the fluorescence light is collected by the objective lens and descanned by the same galvanometer as the illumination beam, purified the in-depth signal with a slit, scanned by another galvanometer, and illuminated the MLA for the light-field imaging.

### Network implementation of AIR

The iterative process of our AIR network can be delineated into four distinct steps: 1) position encoding; 2) INR reconstruction; 3) View-wise aberration prediction; and 4) loss computation and network optimization.

### Position encoding

Deep networks exhibit a predisposition toward learning lower frequency functions. By employing position encoding functions to map 3D coordinates into a higher-dimensional space before they are input into the network, this approach facilitates more precise data fitting processes involved in high-frequency variations[57]. The expression of the Fourier position encoding function employed in this paper is provided below:

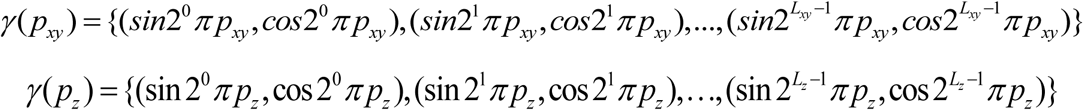

Where *p*_*xy*_represents the lateral coordinates (x, y), while *p*_*z*_ denotes the axial coordinate. The parameters *L*_*xy*_ and *L*_*z*_ correspond to the lateral and axial encoding orders, respectively. Following the position encoding process, a high-dimensional feature vector, denoted as *V* ⊆ *R*^*N*×*L*^ is generated, where N signifies the number of input coordinates and L represents the number of channels of the encoded features determined by *L*_*xy*_ and *L*_*z*_. Each element *V*_*i*_ within the vector V can be expressed as follows:

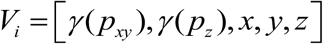

### INR reconstruction

Our INR (Implicit Neural Representation) reconstruction network utilizes a multi-layer perceptron (MLP) structure with ReLU activation. 3D coordinates are firstly flattened into a one-dimensional vector, transforming the spatial structure into a format suitable for the MLP architecture. Next, the 3D coordinates undergo Fourier position encoding, after which they are fed into the INR reconstruction network. The INR reconstruction network predicts the intensity values at each coordinate. This process can be represented as follows:

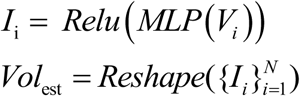

Here, i denotes the index of a specific voxel, and *I*_*i*_ represents the corresponding grayscale value. The ReLU activation function is employed to truncate any predicted values that are less than zero. Following the inference process of the network, the predicted voxel values are subsequently reshaped into a 3D estimated volume (*Vol*_*est*_).

### View-wise aberration prediction

In order to constrain the 3D estimated volume of the INR reconstruction network using the 2D input raw light field images, we must generate a PSF that aligns with the parameters of the optical system to project 3D estimated volume, thereby producing the 2D estimated light field image. Subsequently, the estimated light field image is compared with the input raw light field image to compute a loss function, which serves to constrain the output of the INR reconstruction network. The use of a PSF based on wave optics does not account for aberrations and distortions caused by the sample and optical system, leading to artifacts and ghosting in the reconstructed image (Fig. 1). Addressing this issue, optimizing the Zernike coefficients and using linear combinations of Zernike terms to represent aberrations has been an effective approach for removing aberrations in wide-field imaging[58]. Here, we introduce a view-wise aberration prediction network with learnable parameters 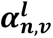 representing Zernike coefficients. Here, v denotes the viewing angle index, ranging from 1 to 9. while n and l represent radial and azimuthal indices, respectively, identifying the specific Zernike term (spherical or coma, etc.). A total of 12 terms are employed (ANSI standards 4th to 15th). Thus, the estimated aberration for each view (𝝓_𝒗_) can be computed as follows:

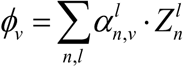

The integration of estimated aberration into the light field PSF is an important problem. Yi Zhang et al. addressed this by introducing optical aberrations on the native image plane and simulating the PSF after passing through a microlens array modulation using Fresnel diffraction[59]. However, as the PSF for Fourier LFM is essentially 5D, it requires substantial computational resources, particularly for deep learning networks where the process needs to be repeated every iteration. Therefore, in this study, we chose to directly input the ideal PSF from wave optics and add aberrations to the PSF in the spatial frequency domain for each view. The resulting aberrated PSF was then inverse Fourier-transformed to yield the final PSF with aberrations 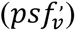. This approach not only saves unnecessary computational resources but also enables the independent addition of aberrations to the PSF for each view (*ps*𝑓_*v*_). The process can be represented as follows:

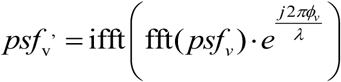

### loss computation and network optimization

Having obtained a PSF with aberrations, forward projection of the 3D estimated volume produced by the INR reconstruction network is performed. This involves a slice-wise convolution of the 3D estimated volume with the estimated PSF along the depth direction, followed by sum projection to generate the 2D estimated light field image (*LF*_*est*_). The error between this estimated light field image and the input raw light field image (*LF*_𝑟𝑎𝑤_) is quantified using a designated loss function:

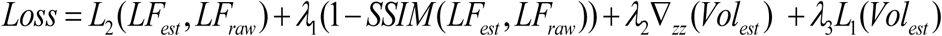

Where *L*_2_ is *L*_2_loss function. SSIM represents the calculation of the structural similarity between two images. ∇_*zz*_ and *L*_1_ are the regularization term to avoid model overfitting. 𝜆_1_, 𝜆_2_, 𝜆_3_ are weighting coefficients.

An Adam optimizer then utilizes the gradients derived from this loss function to jointly update the parameters of both the INR reconstruction network and the view-wise aberration prediction network. This constitutes a single iteration of the AIR network. With increasing iterations, the INR reconstruction network produces a 3D structure of higher fidelity, while the aberration prediction network refines its prediction of the aberrations based on the input raw light field image.

### Synthesis and characterization of TTT8,4-B

TT6,2-B was synthesized as described previously[60]. ^1^H NMR (400 MHz, chloroform-*d*) δ 7.57 (d, *J* = 8.6 Hz, 4H), 7.28 (dd, *J* = 8.5, 7.2 Hz, 10H), 7.17–7.12 (m, 8H), 7.11–7.02 (m, 8H), 2.57 (d, *J* = 7.0 Hz, 4H), 1.39–1.29 (m, 2H), 1.14–0.86 (m, 16H), 0.64 (td, *J* = 7.0, 1.5 Hz, 6H), 0.47 (t, *J* = 7.4 Hz, 6H). Similarly, the synthesis of TT8,4-B was carried out following a reported procedure[61,62]. ^1^H NMR (600 MHz, chloroform-*d*) δ 7.57 (d, *J* = 8.6 Hz, 4H), 7.32–7.21 (m, 10H), 7.18–7.12 (m, 8H), 7.12–7.07 (m, 4H), 7.07–7.03 (m, 4H), 2.58 (d, *J* = 7.0 Hz, 4H), 1.45–1.37 (m, 2H), 1.14 (p, *J* = 7.2 Hz, 4H), 1.07–0.89 (m, 28H), 0.80 (t, *J* = 7.3 Hz, 6H), 0.64 (td, *J* = 7.1, 1.6 Hz, 6H).

#### Synthesis of intermediate 1

A solution of 1-(4-bromophenyl)-1,2,2-triphenylethylene (2.05 g, 5 mmol), aniline (0.6 g, 6.5 mmol), tri-tert-butylphosphine (16.2 mg, 0.08 mmol), Pd_2_(dba)_3_ (64 mg, 0.07 mmol), and sodium tert-butoxide (625 mg, 6.5 mmol) were refluxed under nitrogen (N_2_) in dry toluene (30 mL) at 110 °C for 24 h. After cooling to room temperature, the solvent was removed by evaporation under reduced pressure. H_2_O (30 mL) and chloroform (200 mL) were then added. The organic layer was separated and washed with brine, dried over anhydrous MgSO_4_, and evaporated to dryness under reduced pressure. The crude product was purified by column chromatography on silica gel using hexane/chloroform (v/v = 5/1) as eluent to afford intermediate **1**. ^1^H NMR (600 MHz, chloroform-*d*) δ 7.26 (d, *J* = 7.6 Hz, 2H), 7.23–6.99 (m, 18H), 6.93 (d, *J* = 8.3 Hz, 3H), 6.84 (d, *J* = 8.2 Hz, 2H).

#### Synthesis of intermediate 2

Intermediate **1** (3.77 g, 8.9 mmol), 4-bromoiodobenzene (2.97 g, 10.6 mmol), and 1,10-phenanthroline (0.27 g, 1.5 mmol) were dissolved in toluene (30 mL). After the solution was heated to 100 °C, CuCl (0.15 g, 1.5 mmol) and KOH (1.23 g, 22.0 mmol) were added under N_2_ purge. The mixture was refluxed for 48 h at 120 °C. After cooling to room temperature, the solvent was removed by evaporation under reduced pressure. H_2_O (30 mL) and chloroform (40 × 3 mL) were then added. The organic phase was separated and washed with brine, and dried over anhydrous Na_2_SO_4_. After removal of the solvent, the residual was purified on a silica gel column with ethyl acetate/petroleum (v/v = 1/40) as eluent, giving intermediate **2**. ^1^H NMR (400 MHz, chloroform-*d*) δ 7.32 (d, *J* = 8.9 Hz, 2H), 7.29-7.23 (m, 2H), 7.17-7.02 (m, 18H), 6.91 (dd, *J* = 8.8, 2.9 Hz, 4H), 6.80 (d, *J* = 8.6 Hz, 2H).

#### Synthesis of intermediate 3

Intermediate **2** (2 g, 3.46 mmol), bis(pinacolato)diboron (1.93 g, 7.61 mmol), Pd(dppf)_2_Cl_2_ (119.3 mg, 0.16 mmol), and KOAc (1.02 g, 10.2 mmol) were added into a two-necked flask. Then, 1,4-dioxane was added under an N_2_ atmosphere. The mixture was reacted at 90 °C for 12 h. After being cooled to room temperature, the solvent was evaporated under reduced pressure.Subsequently, 30 mL of H_2_O and 200 mL of chloroform were added. The organic layer was separated and subjected to brine washing, followed by drying with anhydrous MgSO_4_. Finally, the mixture was evaporated to dryness under reduced pressure. The crude product was purified by a silica gel column (petroleum ether/dichloromethane = 3/1, v/v) to give a yellow powder (1.25 g, yield: 61.4%). ^1^H NMR (600 MHz, chloroform-*d*) δ 7.69 (d, *J* = 8.0 Hz, 2H), 7.27 (d, *J* = 7.8 Hz, 2H), 7.21–7.04 (m, 18H), 7.01 (d, *J* = 8.0 Hz, 2H), 6.93 (d, *J* = 8.1 Hz, 2H), 6.86–6.83 (m, 2H), 1.38 (d, *J* = 2.8 Hz, 12H).

#### Synthesis of TTT8,4-B

The synthesis of TTT8,4-B followed the procedure outlined in the previous report.^[2]^ Briefly, intermediate **3** (0.63 g, 1 mmol), BBTD-T(8,4) (0.34 g, 0.4 mmol), Pd_2_(dba)_3_ (22.9 mg, 0.025 mmol), P(o-tol)_3_ (60.9 mg, 0.2 mmol), toluene (1.5 mL), and H_2_O (0.3 mL) were added and sealed in a microwave tube. The mixture was reacted at 120 °C for 12 h. After being cooled to room temperature, the solvent was evaporated under reduced pressure. Subsequently, 30 mL of H_2_O and 60 mL of dichloromethane were added. The organic layer was separated and subjected to brine washing, followed by drying with anhydrous MgSO_4_. Finally, the mixture was evaporated to dryness under reduced pressure. The crude product was purified by a silica gel column (petroleum ether/dichloromethane = 2/1, v/v) to give a green powder (0.388 g, yield: 40%). ^1^H NMR (500 MHz, chloroform-*d*) δ 7.47 (d, *J* = 8.4 Hz, 4H), 7.20 (s, 4H), 7.17 (s, 2H),7.12–6.94 (m, 40H), 6.84 (d, *J* = 7.9 Hz, 4H), 6.77 (s, 4H), 2.50 (d, *J* = 6.8 Hz, 4H), 1.34 (dd, *J* = 12.7, 6.6 Hz, 2H), 1.08 (h, *J* = 7.1 Hz, 4H), 0.95–0.82 (m, 26H), 0.73 (t, *J* = 7.3 Hz, 6H), 0.60–0.54 (m, 6H). ^13^C NMR (126 MHz, dichloromethane-*d*) δ 153.23, 147.42, 147.21, 146.07, 145.64, 144.80, 143.99, 143.82, 143.60, 140.79, 140.69, 138.67, 132.16, 131.27, 131.25, 131.22, 129.28, 128.96, 128.10, 127.63, 127.60, 126.49, 126.42, 126.38, 126.31, 125.32, 124.67, 123.39, 123.29, 116.25, 39.07, 34.76, 33.25, 32.97, 31.80, 29.53, 28.60, 28.57, 26.32, 26.29, 22.82, 22.65, 13.86, 13.80. MS: m/z: [M]^+^ calculated for C_114_H_108_N_6_S_4_: 1689.75911; Found: 1689.75607.

### Preparation of TTT8,4-B NPs and TTT8,4-B NPs-αOX40

For the preparation of TTT8,4-B NPs, 1 mL of THF solution containing 1 mg of TTT8,4-B and 10 mg of PS5000-mPEG5000 was first prepared. This solution was then mixed with 9 mL of ultrapure water under continuous ultrasonication for 2 minutes using a microtip probe sonicator (45% output power). After ultrasound, the reaction mixture was transferred into a dialysis bag (MWCO 3500 Da) and dialyzed against ultrapure water for 24 h, with the water refreshed every 4 h to ensure the complete removal of THF. Following dialysis, the resulting NPs dispersed in ultrapure water were concentrated using an ultrafiltration tube and either stored at 4 °C or used immediately.

To prepare TTT8,4-B NPs-αOX40, maleimide-modified TTT8,4-B NPs (TTT8,4-B NPs-Mal) were first synthesized. The procedure for preparing TTT8,4-B NPs-Mal was identical to that of TTT8,4-B NPs, except that the 1 mL THF solution contained 8 mg of PS5000-mPEG5000, 2 mg of PS5000-PEG5000-Mal, and 1 mg of TTT8,4-B. Subsequently, TTT8,4-B NPs-Mal were functionalized with thiol-modified αOX40 (prepared according to the literature[63,64]) via sulfhydryl-maleimide coupling to yield the final TTT8,4-B NPs-αOX40. The resulting TTT8,4-B NPs-αOX40 in PBS were concentrated and stored at 4 °C or used immediately.

The concentrations of TTT8,4-B NPs and TTT8,4-B NPs-αOX40, based on TTT8,4-B content, were quantified using a pre-established standard absorption curve of TTT8,4-B in THF. More details of the experimental section and characterization can be found in Supplementary Text S2.

### Phantom preparation

The dried leaf was firstly dyed in TTT8,4-B NPs (1.7 mmol/mL) and Rhodamine B (1 mg/mL) mixture for 30 min, which provides fluorescent signals simultaneously in visible and near-infrared. Then, it was fixed with a thin layer of transparent agarose (0.8%). After the acquisition of the surface image without scattering tissue, the lipid (0.167%) and agarose (0.8%) mixture was applied on the top of the phantom. The depth of the covered lipid tissue was estimated by calculating the volume and the area of the disk. Once a new layer of lipid agarose was applied, the phantom was placed on a flat surface and waited for solidification. The photos of the leaf vessels under different depths of lipid can be found in Fig. 2a.

### Hemodynamic imaging

Fig. S42 is a schematic of the process of measuring flow speed with the NIR-II LFM strategy. First, the depth map was calculated by searching through the points with the maximum value in the vessel. Then, a skeleton was drawn on an interested vessel in the MIP images. By combining the depth information and the top view 2D location matrix, we can derive the volumetric skeleton location coordinates. A kymograph was relocated and constructed with the volumetric coordinates. The 3D blood flow velocity of the interested vessel was quantified based on the relationship between the slope with the real speed.

### Animal preparation for *in vivo* brain imaging

We used BALB/C mice (mice, male, 8 weeks old) for the *in vivo* studies. Throughout the experiments, the mice were maintained under anesthesia with 1.5% isoflurane and the mice body were kept at 37 °C using a heating pad. Craniotomy surgery was performed to remove the skull, and a thin glass plate was used to seal the brain. Before imaging, erythromycin cream was applied to mice eyes to keep them moist. The laser power was measured with a power meter and controlled carefully. All experimental procedures were carried out in conformity with the animal protocol approved by the laboratory animal center at Huazhong University of Science and Technology.

### Mouse imaging under norepinephrine

To validate the system performance in monitoring acute hemodynamic change. The mouse was imaged under the challenge of norepinephrine. The mouse was anesthetized and placed on a heating pad. The AIE NP solution was injected before imaging and the region was selected under LFM. A syringe was injected intravenously without drug delivery. After acquiring baseline vasculature and flow speed for 1 min. We pumped norepinephrine (0.02 mg/mL, 200 μL) and monitored for another 1.5 mins.

### Ischemic stroke

We used 532 nm CW laser and a photosensitizer (Rose Bengal, Sigma; concentration 10 mg/mL, 0.2 mL per mouse) to generate blood clots in target regions. Specifically, the mixture of Rose Bengal and TTT8,4-B NPs were injected intravenously. Then, the mouse was placed under the objective. Water was applied to the top of the head and the objective was soaked. The NIR laser was turned on with a safe level output power. The region of interest was located with the aid of NIR-II LFM. Then, the brain region was continuously monitored for 1 min at a 20 Hz frame rate and the 532 nm laser was launched to the target vessel. The 532 nm laser dwelled for 30 seconds for the stable generation of stroke. Finally, the *in vivo* mouse brain was recorded for another minute.

### Tumor inoculation and *in vivo* NIR-II LFM imaging of tumors

5-week-old BALB/c mice were shaved using hair removal lotion and inoculated with ∼ 2 million CT26 cancer cells near the inguinal region for tumor growth. When the tumor size reached ∼ 4 mm, 100 μL aOX40-TTT8,4-B NPs were intravenously injected into the BALB/c mouse through tail vein 24 hours after 50-μg CpG (ODN 2395, InvivoGen) intratumoral treatment. NIR-II LFM was performed to track the 3D movement of aOX40-TTT8,4-B NPs-stained T cells in the CT26 tumor 24 hours post the probe injection.

## List of Abbreviations

(NIR-II): near-infrared II
(LFM): light-field microscopy
(INR): implicit neural representation
(AIR): aberration-corrected implicit neural representation reconstruction
(DAO): digital adaptive optics
(TPE): triphenylethylene
(THF): tetrahydrofuran
(NPs): nanoparticles
(TEM): transmission electron microscopy
(QY): quantum yield
(ICG): indocyanine green
(HOMO): highest occupied molecular orbital
(LUMO): lowest unoccupied molecular orbital
(Rg): gyration
(TICT): twisted intramolecular charge transfer
(FLFM): fourier light field microscopy
(DOF): depth of field
(VAP): view-wise aberration prediction
(MLP): multi-layer perceptron
(SSIM): structure similarity index measurement
(SBR): signal-to-background ratio
(CpG): cytosine-phosphate-guanine
(TLR9): Toll-like receptor 9

## Supplementary information

Supplementary Figs. 1-44, Notes 1-2, Table 1 and Movies 1-6.

## Availability of data and materials

The raw data will be made available upon request to the corresponding author. The remaining data are available within the Article, Supplementary Information.

## Author contributions

D.L. conceived the idea. F.Z. and M.H. led the experiment with supervision from D.L. and P.F.. M.H., F.Z. and Y.H. built the optical setup. Z.Z., X.L. and K.R. designed, synthesized and characterized the TTT8,4-B. Z.Z. and X.L. designed, fabricated, and characterized the TTT8,4-B nanoparticles (NPs) and TTT8,4-B NPs-αOX40. M.H. and F.Z. discussed and designed the experimental system and control system. F.Z., M.H. and S.M. handled the animal preparation. F.Z., M.H., Y.H., C.Y. and X.H. processed and analyzed microscopic data. Z.Z, D.L. and P.F. manage the project. F.Z., X.L., M.H., C.Y., M.K., D.W., Z.Z., D.L. and P.F. wrote and revised the paper.

## Acknowledgements

We thank Yao Zhou from Huazhong University of Science and Technology for her kind advise. We also thank the Research Core Facilities for Life Science (HUST) for the help of two-photon measurement.

## References

1. Hoover, E. E. & Squier, J. A. Advances in multiphoton microscopy technology. Nature Photonics 7, 93–101 (2013).

2. Samanta, S. et al. AIE-active two-photon fluorescent nanoprobe with NIR-II light excitability for highly efficient deep brain vasculature imaging. Theranostics 11, 2137–2148 (2021).

3. Horton, N. G. et al. In vivo three-photon microscopy of subcortical structures within an intact mouse brain. Nature Photonics 7, 205–209 (2013).

4. Wang, L. V. & Yao, J. A practical guide to photoacoustic tomography in the life sciences. Nature Methods 13, 627–638 (2016).

5. Zhong, F., Huang, X., Sun, M., Li, D. & Fei, P. High-Speed Hemodynamic Imaging with Low-Fluence Photoacoustic Microscopy and Self-Supervised Single Volume Denoising. Laser and Photonics Reviews 19, 2401291 (2025).

6. Lan, B. et al. High-speed widefield photoacoustic microscopy of small-animal hemodynamics. Biomedical Optics Express 9, 4689 (2018).

7. Zi-Han, C. et al. NIR-II Anti-Stokes Luminescence Nanocrystals with 1710 nm Excitation for in vivo Bioimaging. Angewandte Chemie International Edition e202416893, 1–7.

8. Xiangdan, M. et al. Recent Advances in Near-Infrared-II Fluorescence Imaging for Deep-Tissue Molecular Analysis and Cancer Diagnosis. Small 18, 1–26.

9. Feng, Z. et al. Engineered NIR-II fluorophores with ultralong-distance molecular packing for high-contrast deep lesion identification. Nature Communications 14, 5017 (2023).

10. Feng, Z. et al. Perfecting and extending the near-infrared imaging window. Light Science Applications 10, 197 (2021).

11. Hong, G. et al. Through-skull fluorescence imaging of the brain in a new near-infrared window. Nature Photonics 8, 723–730 (2014).

12. Qi, J. et al. Aggregation-Induced Emission Luminogen with Near-Infrared-II Excitation and Near-Infrared-I Emission for Ultradeep Intravital Two-Photon Microscopy. ACS Nano 12, 7936–7945 (2018).

13. Welsher, K. et al. A route to brightly fluorescent carbon nanotubes for near-infrared imaging in mice. Nature Nanotechnology 4, 773–780 (2010).

14. Demas, J. et al. High-speed, cortex-wide volumetric recording of neuroactivity at cellular resolution using light beads microscopy. Nature Methods 18, 1103–1111 (2021).

15. Papagiakoumou, E., Ronzitti, E. & Emiliani, V. Scanless two-photon excitation with temporal focusing. Nature Methods 17, 571–581 (2020).

16. Wang, F. et al. Light-sheet microscopy in the near-infrared II window. Nature Methods 16, 545–552 (2019).

17. Wang, F. et al. In vivo NIR-II structured-illumination light-sheet microscopy. Proceedings of the National Academy of Sciences of the United States of America 118, e2023888118 (2021).

18. Wang, Z. et al. Real-time volumetric reconstruction of biological dynamics with light-field microscopy and deep learning. Nature Methods 18, 551–556 (2021).

19. Zhang, Y. et al. Long-term mesoscale imaging of 3D intercellular dynamics across a mammalian organ. Cell 187, 6104–6122.e25 (2024).

20. Yi, C. et al. Video-rate 3D imaging of living cells using Fourier view-channel-depth light field microscopy. Communications Biology 6, 1259 (2023).

21. Zhang, Y. et al. Multi-focus light-field microscopy for high-speed large-volume imaging. PhotoniX 3, 30 (2022).

22. Yoon, Y.-G. et al. Sparse decomposition light-field microscopy for high speed imaging of neuronal activity. Optica 7, 1457 (2020).

23. Truong, T. V. et al. High-contrast, synchronous volumetric imaging with selective volume illumination microscopy. Communications Biology 3, 74 (2020).

24. Hsu, F.-C. et al. Light-field microscopy with temporal focusing multiphoton illumination for scanless volumetric bioimaging. Biomedical Optics Express 13, 6610 (2022).

25. Prevedel, R. et al. Simultaneous whole-animal 3D imaging of neuronal activity using light-field microscopy. Nature Methods 11, 727–730 (2014).

26. Wagner, N. et al. Instantaneous isotropic volumetric imaging of fast biological processes. Nature Methods 16, 497–500 (2019).

27. Campagnola, P. J. High-speed 3D mapping of nonlinear structures. Nature Photonics 14, 527–528 (2020).

28. Gustafsson, M. G. L. et al. Three-Dimensional Resolution Doubling in Wide-Field Fluorescence Microscopy by Structured Illumination. Biophysical Journal 94, 4957–4970 (2008).

29. Gao, B. et al. NIR-II Light Field Microscopy for Through-Skull Hemodynamic Volumetric Imaging in Awake Mice. Laser and Photonics Reviews 19, 2401685 (2025).

30. Booth, M. J. Adaptive optical microscopy: the ongoing quest for a perfect image. Light Science Applications 3, e165–e165 (2014).

31. Tang, J., Germain, R. N. & Cui, M. Superpenetration optical microscopy by iterative multiphoton adaptive compensation technique. Proceedings of the National Academy of Sciences of the United States of America 109, 8434–8439 (2012).

32. Wang, C. et al. Multiplexed aberration measurement for deep tissue imaging in vivo. Nature Methods 11, 1037–1040 (2014).

33. Wu, J. et al. Iterative tomography with digital adaptive optics permits hour-long intravital observation of 3D subcellular dynamics at millisecond scale. Cell 184, 3318–3332.e17 (2021).

34. Sun, Z. et al. CXCL9+ Macrophage-targeted NIR-II aggregation-induced emission nanoprobes for the early diagnosis of myocarditis. Nano Today 54, 102107 (2024).

35. Yang, S. et al. More Is Better: Dual-Acceptor Engineering for Constructing Second Near-Infrared Aggregation-Induced Emission Luminogens to Boost Multimodal Phototheranostics. Journal of the American Chemical Society 145, 22776–22787 (2023).

36. Zhang, Z. et al. The fast-growing field of photo-driven theranostics based on aggregation-induced emission. Chemical Society Reviews 51, 1983–2030 (2022).

37. Li, X. et al. Aggregation-induced emission materials: a platform for diverse energy transformation and applications. Journal of Materials Chemistry A 11, 4850–4875 (2023).

38. Cai, Y., et al. Near-infrared fluorophores with absolute aggregation-caused quenching and negligible fluorescence re-illumination for in vivo bioimaging of nanocarriers. Aggregate 4, e277 (2023).

39. Li, Y. et al. Design of AIEgens for near-infrared IIb imaging through structural modulation at molecular and morphological levels. Nature Communications 11, 1255 (2020).

40. Lin, R. et al. Type I Photosensitization with Strong Hydroxyl Radical Generation in NIR Dye Boosted by Vigorous Intramolecular Motions for Synergistic Therapy. Advanced Materials 35, 2303212 (2023).

41. Ren, K. et al. The Longer, the Stronger: Alkyl Chain Engineering for High-Brightness NIR-II AIE Nanoprobes towards High-Resolution In Vivo Bioimaging. Preprint at 10.26434/chemrxiv-2025-bctb9-v2 (2025).

42. Grabowski, Z. R., Rotkiewicz, K. & Rettig, W. Structural Changes Accompanying Intramolecular Electron Transfer: Focus on Twisted Intramolecular Charge-Transfer States and Structures. Chemical Reviews 103, 3899–4032 (2003).

43. Gottschalk, S. et al. Rapid volumetric optoacoustic imaging of neural dynamics across the mouse brain. Nature Biomed Engineering 3, 392–401 (2019).

44. Zhang, Z. et al. Imaging volumetric dynamics at high speed in mouse and zebrafish brain with confocal light field microscopy. Nature Biotechnology 39, 74–83 (2021).

45. Kamshilin, A. A. et al. Advancing intraoperative cerebral blood flow monitoring: integrating imaging photoplethysmography and laser speckle contrast imaging in neurosurgery. Frontiers of Optoelectronics 18, 20 (2025).

46. Chen, W. et al. High-throughput volumetric mapping of synaptic transmission. Nature Methods 21, 1298–1305 (2024).

47. Zhong, F. & Hu, S. Thin-film optical-acoustic combiner enables high-speed wide-field multi-parametric photoacoustic microscopy in reflection mode. Optics Letters 48, 195 (2023).

48. Broxton, M. et al. Wave optics theory and 3-D deconvolution for the light field microscope. Optics Express 21, 25418 (2013).

49. Zhao, Z. et al. Two-photon synthetic aperture microscopy for minimally invasive fast 3D imaging of native subcellular behaviors in deep tissue. Cell 186, 2475–2491.e22 (2023).

50. Yao, J. et al. High-speed label-free functional photoacoustic microscopy of mouse brain in action. Nature Methods 12, 407–410 (2015).

51. Wang, X. et al. An Emerging Toolkit of Ho^3+^ Sensitized Lanthanide Nanocrystals with NIR-II Excitation and Emission for *in Vivo* Bioimaging. Journal of the American Chemical Society 147, 2182–2192 (2025).

52. Shih, A. Y. et al. Two-Photon Microscopy as a Tool to Study Blood Flow and Neurovascular Coupling in the Rodent Brain. Journal of Cerebral Blood Flow & Metabolism 32, 33 (2012).

53. Kawarada, Y. et al. Peritumoral CpG DNA Elicits a Coordinated Response of CD8 T Cells and Innate E ectors to Cure Established Tumors in a Murine Colon Carcinoma Model. The Journal of Immunology 167, 5247–5253 (2001).

54. Wagner, H. The immunogenicity of CpG-antigen conjugates. Advanced Drug Delivery Reviews 61, 243–247 (2009).

55. Ruby, C. E., Montler, R., Zheng, R., Shu, S. & Weinberg, A. D. IL-12 Is Required for Anti-OX40-Mediated CD4 T Cell Survival. The Journal of Immunology 180, 2140–2148 (2008).

56. Curti, B. D. et al. OX40 Is a Potent Immune-Stimulating Target in Late-Stage Cancer Patients. Cancer Research 73, 7189–7198 (2013).

57. Rahaman, N., et al. On the Spectral Bias of Neural Networks. International Conference on Machine Learning 97, (2019).

58. Kang, I., Zhang, Q., Yu, S. X. & Ji, N. Coordinate-based neural representations for computational adaptive optics in widefield microscopy. Nature Machine Intelligence 6, 714–725 (2024).

59. Zhang, Y. et al. Computational optical sectioning with an incoherent multiscale scattering model for light-field microscopy. Nature Communications 12, 6391 (2021).

60. Shen, H. et al. Rational Design of NIR-II AIEgens with Ultrahigh Quantum Yields for Photo- and Chemiluminescence Imaging. Journal of the American Chemical Society 144, 15391–15402 (2022).

61. Xu, H., Yuan, L., Shi, Q., Tian, Y. & Hu, F. Ultrabright NIR-II Nanoprobe for Image-Guided Accurate Resection of Tiny Metastatic Lesions. Nano Letter 24, 1367–1375 (2024).

62. Song, S. et al. Molecular engineering of AIE luminogens for NIR-II/IIb bioimaging and surgical navigation of lymph nodes. Matter 5, 2847–2863 (2022).

63. Howard, M. D. et al. Targeting to Endothelial Cells Augments the Protective Effect of Novel Dual Bioactive Antioxidant/Anti-Inflammatory Nanoparticles. Molecular Pharmaceutics 11, 2262–2270 (2014).

64. Parhiz, H. et al. PECAM-1 directed re-targeting of exogenous mRNA providing two orders of magnitude enhancement of vascular delivery and expression in lungs independent of apolipoprotein E-mediated uptake. Journal of Controlled Release 291, 106–115 (2018).

